# Large herbivores link plant phenology and abundance in arctic tundra

**DOI:** 10.1101/2024.02.23.581818

**Authors:** Eric Post, R. Conor Higgins, Pernille Sporon Bøving, Christian John, Mason Post, Jeff Kerby

## Abstract

Plant phenology has been well studied in relation to abiotic conditions and climate change, but poorly studied in relation to herbivory. In contrast, plant abundance dynamics have been well studied in relation to abiotic conditions and herbivory, but poorly studied in relation to phenology. Consequently, the contribution of herbivory to plant phenological dynamics and therefrom to plant abundance dynamics remains obscure. We conducted a nine-year herbivore exclusion experiment to investigate whether herbivory might link plant phenological and abundance dynamics in arctic tundra. From 2009 - 2017 we monitored annual green-up timing and abundance of nine plant taxa, including deciduous shrubs, forbs, and graminoids, on plots that were either grazed or experimentally exclosed from herbivory by caribou (*Rangifer tarandus*) and muskoxen (*Ovibos moschatus*). In 62% of cases, green-up occurred earlier under herbivory, and in 75% of cases abundance was greater under herbivory, compared to green-up and abundance under herbivore exclusion. Moreover, taxa that responded to herbivory with earlier green-up also had comparatively greater abundance later in the growing season. Conversely, taxa that responded to herbivory with delayed green-up exhibited comparatively lower abundance later in the growing season. Hence, well-documented influences of large herbivores on plant abundance and community composition in arctic tundra may, at least to some extent, relate to influences of herbivory on plant phenology. We recommend that ongoing and future assessments of the contribution of herbivores to plant abundance and community responses to climate change, especially in the Arctic, should also consider impacts of herbivores on plant phenology.

## Introduction

Life history theory predicts that plant performance, including annual growth, abundance, and offspring production, should reflect strategies aimed at optimizing the timing of critical life cycle stages such as the timing of growth onset (1). This is because plants risk tissue losses to both adverse abiotic conditions and herbivory, both of which may subsequently influence abundance, survival, and reproductive success (2). Consequently, both the seasonal timing of growth onset and variation in it (i.e., phenology) and abundance of plants should be sensitive to abiotic conditions and herbivory (3). Yet plant phenology is studied nearly exclusively in relation to abiotic conditions (4) while plant abundance, diversity, and community dynamics are well studied in relation to both herbivory (5, 6) and weather or climate (7, 8).

Reflecting this disparity, plant phenology has been remarkably well researched in relation to climate change, representing one of the most commonly focused upon biological responses to recent warming (4, 9–11). This is especially evident in the Arctic (12, 13), where abiotic constraints on the seasonal timing of plant growth are strong (14, 15) and where warming has been rapid and pronounced (16, 17). By comparison, little focus has been placed on plant phenological responses to herbivory (18). Some evidence indicates that large herbivores such as bison (*Bison bison*) can advance plant green-up dynamics by reducing light competition through tissue removal and via promotion of photosynthetic activity by increasing nitrogen content in surviving plant leaf tissue (19). Despite extensive focus on plant phenological responses to warming in the Arctic (20), only two studies have examined impacts of herbivory on plant phenology in the region (21, 22). One of those studies, conducted in a high arctic fen community, revealed that exclusion of muskoxen (*Ovibos moschatus*) advanced the annual timing of peak vegetation biomass (21). The other, conducted in a low arctic shrub-graminoid-forb complex, revealed that exclusion of caribou (*Rangifer tarandus*) and muskoxen delayed the annual onset of plant green-up timing, implying that herbivory accelerated green-up (22). Consequently, much remains to be understood about the role of herbivory in plant phenological dynamics in the Arctic or elsewhere.

This knowledge gap is notable considering large herbivores, including caribou or reindeer (also *R. tarandus*) and muskoxen in the Arctic, have well-documented effects on plant abundance, diversity, and community composition (23–25). Hence, in the absence of better understanding of herbivore impacts on plant phenology we currently lack insights into whether or how such impacts might contribute to well-known effects on plant abundance. As well, the lack of studies linking herbivore impacts on plant phenology to herbivore impacts on plant abundance may reflect an assumption that phenological and abundance dynamics are primarily constrained by different factors (26). Addressing this challenge may be especially important in the context of climate change impacts in the far north because the annual timing of plant phenological events is advancing faster in the Arctic than anywhere else on Earth (27–29) and vegetation in the arctic tundra biome is undergoing complex shifts in composition and structure (30). The extent to which these phenomena may be associated, and the potential contribution of herbivory to any such association, is currently unknown but may be informed by studies such as the one presented here.

## Results and Discussion

We monitored mean annual green-up dates and annual peak abundances of nine arctic tundra plant taxa from 2009 through 2017 at a study site near Kangerlussuaq, Greenland, on plots inside fenced exclosures designed to prevent herbivory by muskox and caribou and on plots on adjacent, unfenced control areas exposed to herbivory (See Methods). Comparison of taxon-specific mean annual green-up dates between exclosed and grazed plots (Figure 1a, small symbols) resulted in calculation of 81 experimental responses ratios quantifying the effect of the herbivore exclusion treatment on green-up timing. Of these 81 response ratios, 79 were non-zero (Table S1a). The two zero values resulted from the fact that, in one year of the experiment, mean green-up dates did not differ between exclosed and grazed plots for the forbs *Draba nivalis* and *Stellaria longipes*. Of the non-zero green-up response ratios, 62% were positive and 38% were negative, indicating that in most cases green-up timing occurred earlier under herbivory than under herbivore exclusion (one-sample binomial test Chi-square = −2.03, *P* = 0.043). Comparison of taxon-specific mean annual abundances on exclosed and grazed plots (Figure 1b,c; small symbols) resulted in 62 experimental response ratios quantifying the effect of the herbivore exclusion treatment on abundance. Of these, 60 were non-zero (Table S1b). There were fewer abundance response ratios than green-up response ratios due to instances in which taxa were detected on control plots but not treatment plots (14 cases), detected on treatment plots but not control plots (1 case), or not detected on control and treatment plots (4 cases) (reported as “missing” in Table S1b). These disparities might have reflected differential influences of herbivory and herbivore exclusion on abundances of rare vs. common taxa over the course of the experiment (31). The two zero-value abundance response ratios indicate that, in a single year, mean abundances of the forbs *Bistorta viviparum* and *Draba nivalis* did not differ between exclosed and grazed plots in those cases. Among the 60 non-zero abundance response ratios, 75% were negative and 25% were positive, indicating that in most cases taxon-specific abundance was greater under herbivory than under herbivore exclusion (one-sample binomial test Chi-square = −3.74, *P* < 0.001).

**Figure 1.**
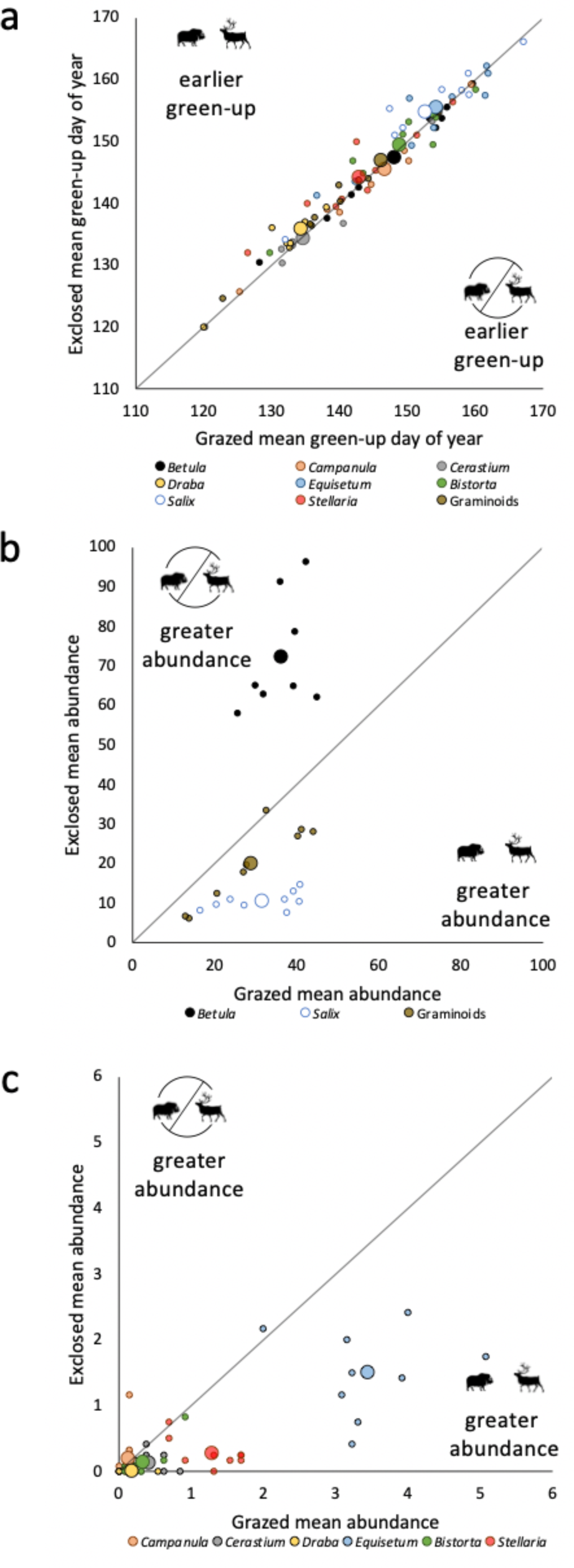
**a)** Mean annual (for each year from 2009 through 2017; small symbols) and pooled mean (across the entire period 2009-17; large symbols) green-up timing by 9 plant taxa at the study site near Kangerlussuaq, Greenland on plots experimentally exclosed from herbivory by caribou and muskoxen plotted against those means on grazed (control) plots. Values above the diagonal parity line are those for which green-up occurred comparatively earlier under exposure to herbivory and values below the diagonal parity line are those for which green-up occurred comparatively earlier under herbivore exclusion. **b and c)** Mean annual (for each year from 2009 through 2017; small symbols) and pooled mean (across the entire period 2009-17; large symbols) abundance (point frame pin intercepts / 0.25m^2^) of 9 plant taxa at the Kangerlussuaq study site on plots experimentally exclosed from herbivory by caribou and muskoxen plotted against those means on grazed (control) plots. Values above the diagonal parity line are those for which abundance was comparatively greater under herbivore exclusion and values below the diagonal parity line are those for which abundance was comparatively greater under exposure to herbivory.

The distributions of positive, negative, or zero-value response ratios varied significantly among the nine focal tundra plant taxa (see Methods) for both green-up timing (Likelihood ratio Chi-square = 26.11, df = 16, *P* = 0.05) and abundance (Likelihood ratio Chi-square = 45.2, df = 16, *P* < 0.001). Two species, the deciduous shrub *Betula nana* and the forb *Bistorta viviparum*, displayed mostly negative green-up response ratios (Table S1a), indicating that green-up by these two species occurred later under herbivory than under herbivore exclusion (Figure 1a, small symbols). In contrast, three taxa, the forb *Draba nivalis*, the deciduous shrub *Salix glauca*, and graminoids, displayed mostly positive green-up response ratios (Table S1a), indicating that green-up by them generally occurred earlier under herbivory than under herbivore exclusion (Figure 1a, small symbols).

The shrub *Betula nana* was the only taxon displaying mostly, and in fact exclusively, positive abundance response ratios (Table S1b), indicating that its abundance was greater under herbivore exclusion than under herbivory (Figure 1b, small symbols). Notably, there was no difference in baseline mean abundance of this species on grazed vs. exclosed plots at the beginning of the experiment (Supplemental Figure S2), indicating that the differences reported here developed over the course of herbivore exclusion as the experiment progressed. In contrast, six taxa, including the forbs *Bistorta viviparum*, *Cerastium alpinum*, *Equisetum arvense*, and *Stellaria longipes*; the shrub *Salix glauca*; and graminoids all displayed mostly negative abundance response ratios (Table S1b), indicating that for these six taxa abundance was comparatively greater under herbivory than under herbivore exclusion (Figures 1b and 1c, small symbols). The greater abundance of *Betula nana* under herbivore exclusion than under herbivory may have contributed indirectly to lower abundances of these six taxa on exclosed plots through competition for space, light, or soil nutrients (e.g., refs 32-34).

Annual abundance response ratios were weakly negatively related to annual green-up response ratios (linear model *R*^2^ = 0.08, *b* = −14.5 ± 6.50, *t* = −2.23, *P* = 0.03) (Figure 2a). Variation in the magnitude of abundance response ratios pooled across the experimental period (2009–17) was more strongly and negatively related to variation in pooled green-up response ratios (Figure 2b). Across taxa, 55% of the variation in the magnitude of tundra plant abundance responses to herbivore removal was explained by responses of their green-up timing to herbivore removal (linear model *R*^2^ = 0.55, *b* = −104.9 ± 35.8, *t* = −2.93, *P* = 0.02). This latter relationship indicates that, over the course of the nine-year experiment, earlier green-up timing under herbivory by six taxa (graminoids; the forbs *Bistorta viviparum*, *Equisetum arvense*, *Stellaria longipes*, *Draba nivalis*; and the shrub *Salix glauca*) was associated with greater abundance under herbivory than under herbivore exclusion later in the growing season (Figure 2b). In contrast, later green-up under herbivory by two taxa (the shrub *Betula nana* and the forb *Campanula gieseckiana*) was associated with lower abundance under herbivory than under herbivore exclusion later in the growing season (Figure 2b).

**Figure 2.**
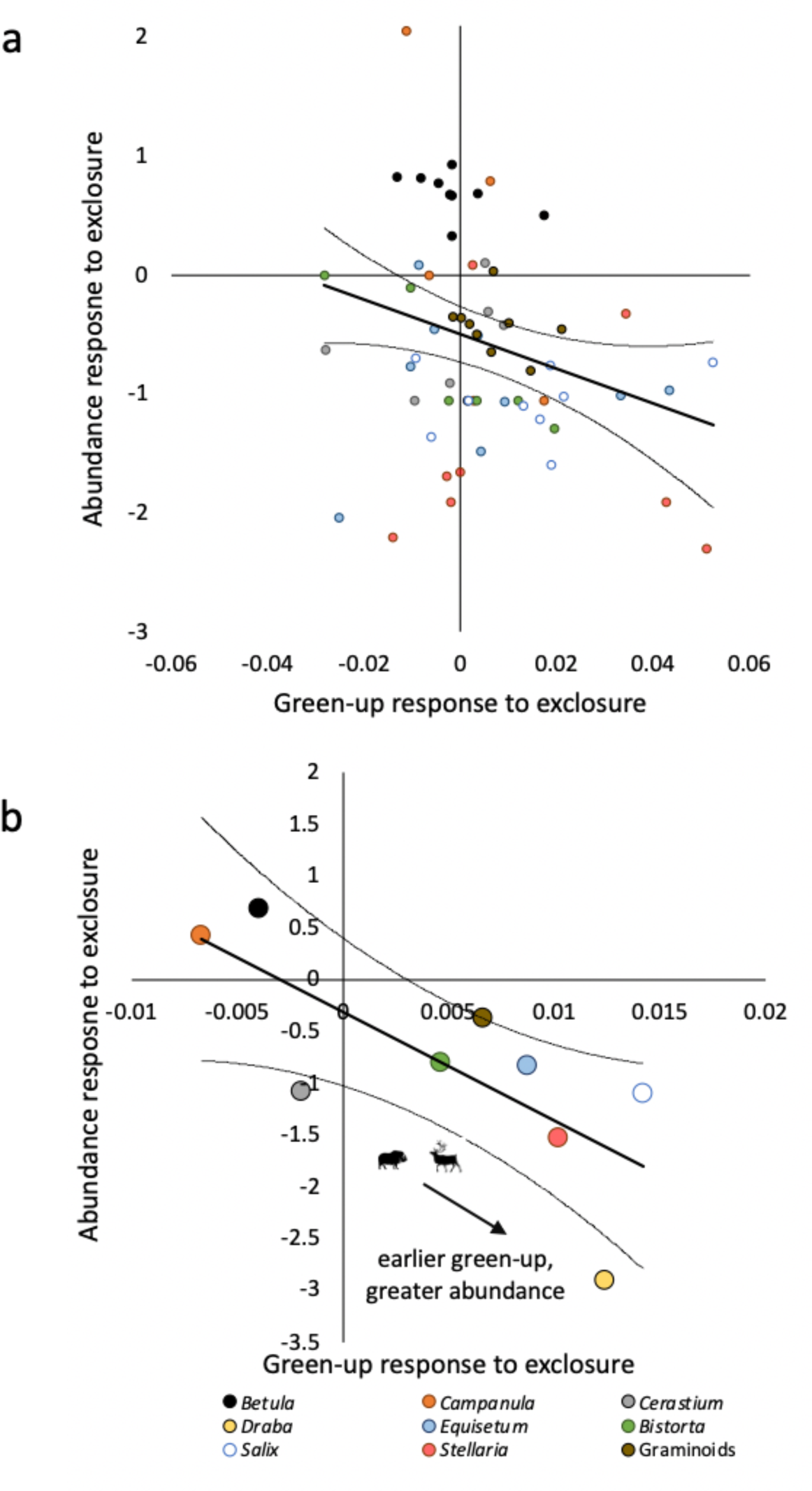
The magnitude of the response of plant abundance to experimental herbivore exclusion in relation to the magnitude of the response of plant green-up timing to experimental herbivore exclusion. Shown are experimental response ratios (see Methods) derived for each of 9 plant taxa using **a)** annual means and **b)** means for data pooled across the experimental period (2009–17).

In general, plant responses to herbivory, or the risk of it, may be expected to take two forms: short-term physiological responses such as the mobilization of resources to increase growth rate, or longer-term adaptive responses to avoid or minimize loss of tissues early in the life cycle that are critical to productivity later in the life cycle (1). This framework thus predicts that plant growth or abundance should be responsive to strategies employed by plants to cope with, or minimize adverse effects of, herbivory through their growth phenology. Our results appear to provide support for this expectation, but also highlight intriguing taxonomic variation in the nature of such responses that warrants further study. For instance, linked plant phenological and abundance responses to herbivory (or its experimental exclusion) did not, in this study, relate clearly to plant growth form or functional group. Notably, the two deciduous shrub species, *Betula nana* and *Salix glauca*, displayed opposing green-up and abundance responses to herbivory (Figure 2b). Similarly, responses of low-growing forbs were mixed, with two species (*Cerastium alpinum* and *Campanula gieseckiana*) exhibiting comparatively later green-up and either lower or greater abundance under herbivory, and four species (*Bistorta viviparum*, *Equisetum arvense*, *Stellaria longipes*, and *Draba nivalis*) all displaying comparatively earlier green-up and greater abundance under herbivory.

Hence, both green-up and abundance responses to our experiment were highly variable across tundra plant growth form and functional group. Intriguingly, however, additional evidence suggests that the relative order in which taxa initiate annual growth in this community may contribute to variation in their green-up responses to herbivory or the risk of it. Taxon-specific green-up responses to herbivory were associated non-linearly with the rank order in which taxa initiate annual growth onset at the study site (Figure S3). Although this relationship is not significant (*P* = 0.16), it suggests that taxa which responded to herbivory with advanced green-up were generally early- and late emerging taxa (Figure S3). This pattern would appear to be consistent with a strategy of minimizing risk of tissue loss under herbivory by those species most likely to experience greatest exposure to herbivory at the tail ends of the annual progression of green-up across the taxa comprising this community (35). In contrast, taxa for which green-up was comparatively delayed by herbivory were those that initiated growth near the middle of the sequence of emergence from the first to last taxon annually (Figure S3).

Attempts to link changes in plant demography, including abundance, to drivers of phenological dynamics have been generally inconclusive (26, 36, 37). This may be at least somewhat attributable to the likelihood that drivers of variation in plant abundance may differ in the magnitude and direction of their effects from drivers of variation in plant phenology. For instance, in arctic tundra plant communities, warming commonly elicits earlier spring growth (38–42), but it may increase or reduce abundance (20, 43–45). Similarly, herbivory itself can increase or reduce plant abundance depending to some extent on plant growth form (2, 23, 25, 32, 46, 47). Our results indicate that even minor effects of herbivore removal on plant emergence phenology may contribute to pronounced effects of herbivore removal on abundance. Moreover, the nature of this relationship suggests that advancement of emergence timing under exposure to herbivory translates into greater relative abundance later in the growing season, consistent with predictions of the induced response framework (48). In contrast, delayed emergence timing in response to herbivory apparently contributes to reduced relative abundance, possibly as a consequence of increased competition for limiting resources (49). Relatedly, a consideration that our simple experimental approach lacks the power to resolve is whether effects of herbivory on phenology are indirect and mediated by effects on abundance. For instance, dwarf birch is the most common species on our plots (31) and the reduction or constraint of its abundance by herbivory may indirectly facilitate phenological escape (i.e., earlier green-up) by under-canopy forbs and graminoids (50, 51). Additional, possibly more complex, experimentation may be needed to address that hypothesis. Nonetheless, our results suggest that the contribution of green-up timing or emergence phenology to plant abundance and community compositional dynamics in arctic tundra warrants further study, especially in the context of interactions between herbivory and climate change.

## Materials and Methods

### Study site, experimental design, and plant phenology and abundance monitoring

The study site is located approximately 20 km east of the village of Kangerlussuaq, Greenland. In June 2002 we initiated a long-term experiment designed to investigate effects of herbivory by caribou and muskoxen on tundra plant abundance, diversity, and community composition. We erected three circular, 800 m^2^ exclosures constructed of woven wire fencing and steel t-posts to exclude caribou and muskoxen. Adjacent to each exclosure we identified a paired control area that remained unfenced and thus open to herbivore access and grazing. The three pairs of exclosure/grazed controls were separated by several hundred meters.

In May 2003, we initiated annual monitoring of taxon-specific plant phenological dynamics inside all three exclosures by recording phenophases of all vascular plants present on four permanently-marked plots in each of the three exclosures (n = 12 herbivore-exclosed plots) on a daily or near-daily basis. This effort resulted in yearly records of annual timing of growth initiation (i.e., basal tissue greening in graminoids, leaf emergence in forbs, and leaf bud opening in deciduous shrubs; hereafter “green-up”), flower set, and blooming for nine vascular plant taxa under herbivore exclusion: dwarf birch (*Betula nana*), gray willow (*Salix glauca*), draba (*Draba* sp.), harebell (*Campanula gieseckiana*), horsetail (*Equisetum arvense*), longstalk starwort (*Stellaria longipes*), alpine chickweed (*Cerastium alpinum*), alpine bistort (*Bistorta vivipara*), and graminoids (sedges and grasses) (22). In May 2009, we added five permanently-marked plots outside of each of the three exclosures, on the adjacent grazed control areas, and initiated monitoring of the same phenophases of all vascular plants occurring on those plots (n = 15 grazed plots). This effort resulted in annual records of seasonal green-up, flower set, and blooming for the same nine taxa on plots exposed to herbivory. Phenological observations commenced in early to mid-May each year and progressed through late June annually (Figure S1). The period of observation was determined by our initial interest in monitoring plant phenology during the annual period when female caribou were present at the site for calving (52) and remained consistent as our research interests expanded (Figure S1). Although muskoxen are present in the area year-round, their seasonal occurrence at the site begins to peak around early May (22). Female caribou migrate into the area annually in early May for calving in late May through early June, before migrating out again in late June (53). The exclosures were removed at the end of the growing season in 2017. Data used here span 2009 through 2017, the period of overlap of phenological monitoring on exclosed and grazed plots. Because we were interested in assessing the role of herbivory in linking vascular plant green-up timing to abundance, we focused our analyses on the subset of phenology data comprising taxon-specific green-up dates.

Annual peak growing season abundances of the same nine taxa were assessed in late July or early August each year (Figure S1) using non-destructive point-frame sampling on a separate but proximal set of permanently marked plots inside (n = 12) and outside (n = 13) of the same exclosures from 2009 through 2017. Point-frame sampling was conducted using a 0.25 m^2^ clear Plexiglas tabletop frame on four adjustable legs. A consistent orientation of the frame during each sampling event was ensured by anchoring the legs in four hollow aluminum pegs set into the ground at the corners of each plot at the cardinal directions. The Plexiglas tabletop frame was drilled with 20, randomly located holes. During sampling, a steel welding pin was lowered through each hole in succession and each contact of the tip of the pin was recorded to species, genus, or functional group according to the nine taxa listed above. We recorded phenological observations on plots separate from those on which abundances were recorded to avoid the potential, however unlikely, for altering phenology through contact with plants (54).

### Analytical approach

Data on both green-up timing (22) and annual abundances (55–57) of the focal taxa have been analyzed previously in relation to the exclosure treatment. Here, we used both sets of data to assess whether the abundance responses of the focal taxa to the exclosure treatment scale with their green-up responses to that treatment. To achieve this, we calculated experimental response ratios of both the green-up response and the abundance response of each taxon to the exclosure treatment both annually for each year from 2009 through 2017 and over the pooled period 2009 - 17. This was done using the following formula:

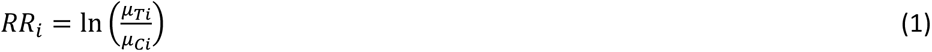

In which *RR_i_* is the experimental response ratio for taxon *i*, *µ_Ti_* is the treatment mean green-up date or abundance for taxon *i* for a given year in the case of annual response ratios or for the period 2009 - 17 in the case of pooled response ratios, and *µ_Ci_* is the control mean green-up date or abundance for taxon *i* for a given year in the case of annual response ratios or for the period 2009 - 17 in the case of pooled response ratios. Thus, *RR* < 0 when green-up is later on grazed compared to exclosed plots or when abundance is greater on grazed than on exclosed plots. To test whether plant abundance responses to herbivory were related to their phenological responses to herbivory, we analyzed variation in taxon-specific abundance response ratios as a function of variation among taxon-specific green-up response ratios. We did this both for taxon-specific response ratios calculated using annual means as well as for taxon-specific response ratios calculated using means pooled for 2009 - 17. In each approach, we used a simple linear model with the former as the dependent variable and the latter as the predictor variable. This was intended to determine whether, and in which direction, the abundance responses of the focal taxa to herbivore removal scaled with their green-up responses to herbivore removal. Although herbivore exclusion is the experimental treatment in this study, we describe the results of the response ratio analyses largely in terms of responses to herbivory. We employ this approach to lend interpretability to our results in the context of the contextual framework described in the Introduction.

## Author Contributions

Conceptualization: EP, RCH, PSB, CJ, MP, JK

Methodology: EP, PSB, JK

Analysis: EP, RCH

Investigation: EP, RCH, PSB, CJ, MP, JK

Data Curation: EP, RCH, JK

Writing - Original Draft Preparation: EP

Writing - Review & Editing: EP, RCH, PSB, CJ, MP, JK

Visualization: EP, RCH, PSB

Supervision: EP

Project Administration: EP

Funding Acquisition: EP, JK

## Acknowledgments

We thank Mads Forchhammer for critical input on experimental design and inspiration for this study, the staff at Kangerlussuaq International Science Support for assistance with project development and logistical arrangements, and Ben Lee for constructive comments on the manuscript. We gratefully acknowledge support from the U.S National Science Foundation (NSF) under grants <u>0124031</u>, <u>0217259</u>, <u>0732168</u>, <u>0713994</u>, <u>1107381</u>, and <u>1525636</u>, and the National Geographic Society to E.P.; and from the European Union’s Horizon 2020 research and innovation program under the Marie Skłodowska-Curie grant (754513), and the Aarhus University Research Foundation to J.K.

## Data availability

Data on plant green-up dates and abundances are available at the Arctic Data Center (58, 59).

## Supplementary Information

**Fig. S1.**
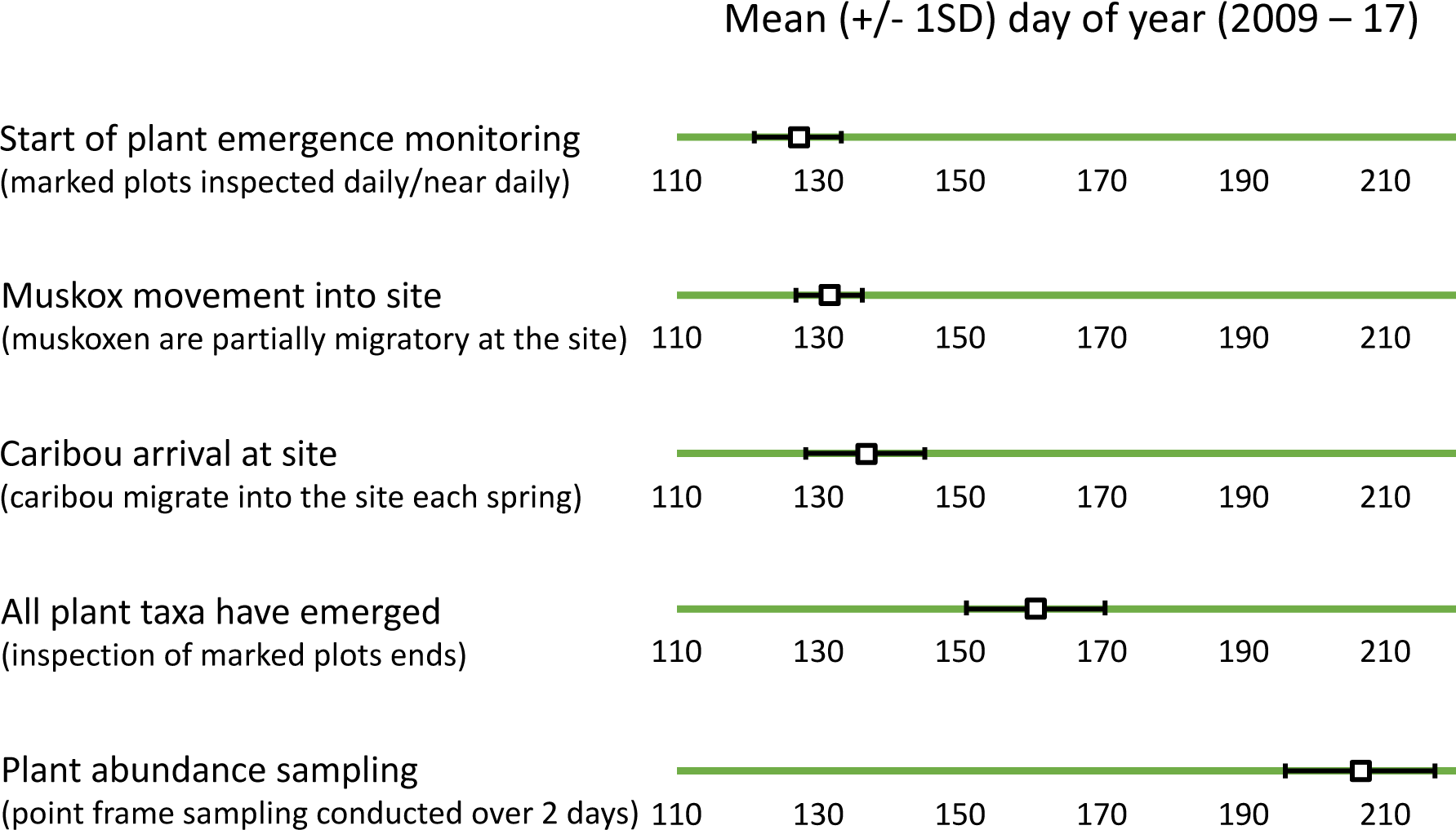
Mean (± 1 standard deviation) day of year on which: observations of plant phenology commenced; muskoxen began to take up residence within the study site, estimated as the date on which 5% of the cumulative total number of muskoxen observed annually were counted; caribou began to migrate into the study site, estimated as the date on which 5% of the cumulative total number of caribou observed annually were counted; all plant taxa occurring on plots monitored for phenology had been recorded as emergent; and non-destructive plant abundance sampling was conducted for the period 2009 through 2017 at the study site near Kangerlussuaq, Greenland (1–3).

**Fig. S2.**
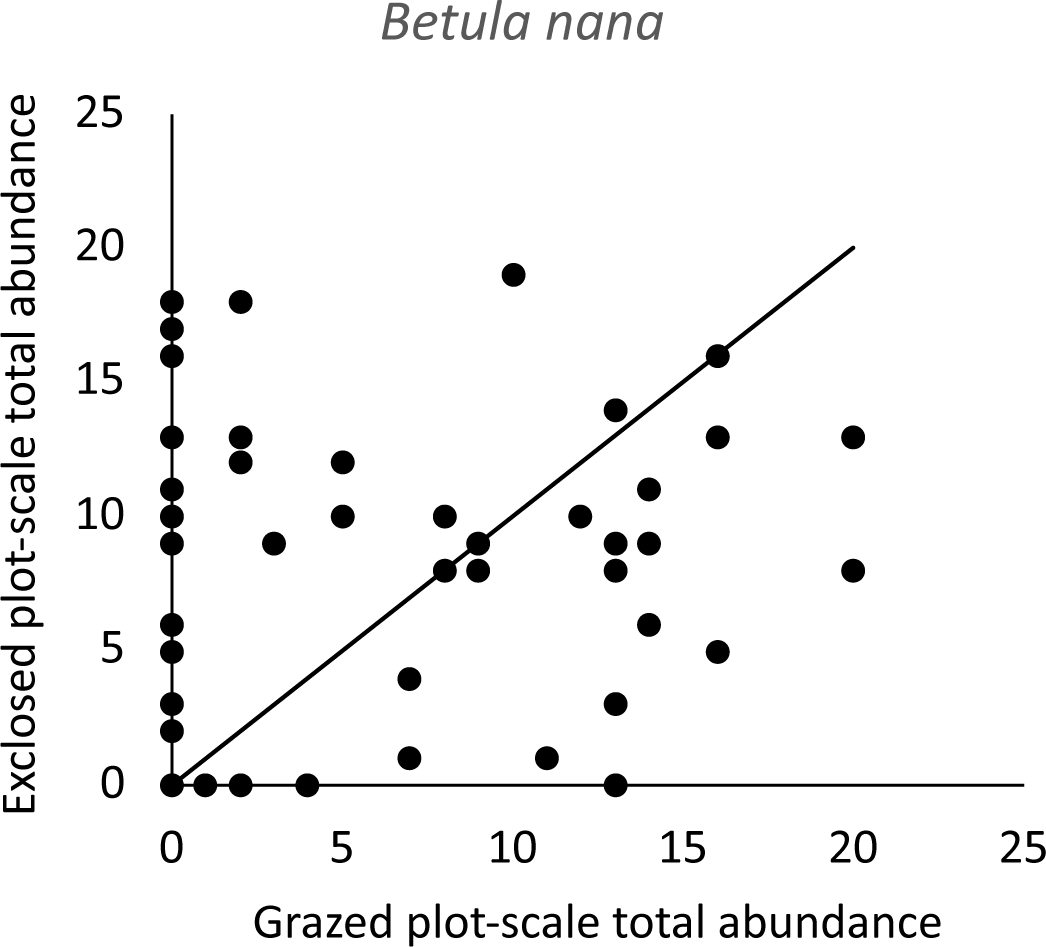
Baseline abundance of *Betula nana* on exclosed and grazed plots assessed in 2002 at the initiation of the large herbivore exclosure treatment at the study site near Kangerlussuaq, Greenland (4). Shown are the total number of pin intercepts counted per plot using a linear, 1m point frame with 10 pin drops spaced at 10cm intervals. Data are displayed as total plot-scale intercepts on individual treatment plots inside exclosures vs intercepts on correspondingly paired individual control plots outside exclosures pooled across the three paired exclosures and grazed controls. A generalized linear model with abundance as the response variable, treatment and exclosure identity as factors, and an identity link function revealed no difference between mean exclosed and grazed abundance (Wald Chi-square = 0.96, df = 1, *P* = 0.33).

**Fig. S3.**
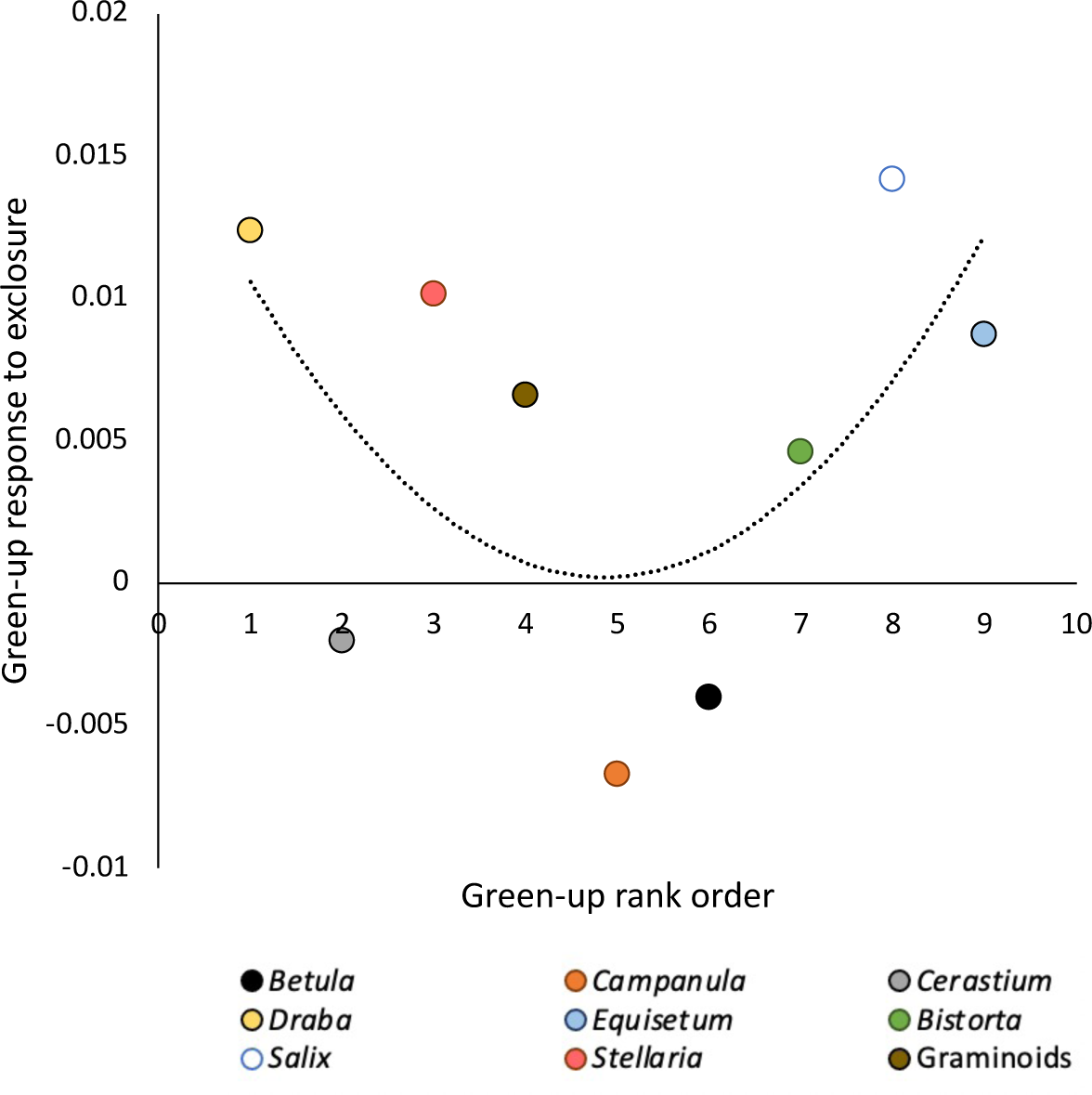
Association between taxon-specific response of plant green-up timing to experimental herbivore exclusion and the rank order in which taxa initiated green-up at the study site near Kangerlussuaq, Greenland over the study period (2009–17). Y-axis values are green-up response ratios shown in Figure 2b in the main text. X-axis values were derived by ranking from 1 (first) through 9 (last) mean dates of green-up for each taxon on grazed plots for the period 2009 - 17 (i.e., the X-axis values for each taxon in Figure 1a in the main text). The dashed line shows the fit of the non-linear model Y = aX^2^ + bX + c (*R*^2^ = 0.34, *F*_3,6_ = 2.49, *P* = 0.16).

**Table S1.** Counts of negative, positive, and zero-value response ratio across nine taxonomic groups for (a) annual mean green-up timing and (b) annual mean abundance responses to the herbivore exclosure treatment from 2009 through 2017 at the study site near Kangerlussuaq, Greenland.

|  |  |  |  |  |  |  |  |  |  |  |  |
| --- | --- | --- | --- | --- | --- | --- | --- | --- | --- | --- | --- |
| **a** |  |  |  |  |  |  |  |  |  |  |  |
| Count |  |  |  |  |  |  |  |  |  |  |  |
|  |  | Taxon |  |  |  |  |  |  |  |  |  |
|  |  | Betula | Bistorta | Campanula | Cerastium | Draba | Equisetum | Graminoids | Salix | Stellaria | Total |
| Green-up response ratio category | negative | 7 | 3 | 5 | 4 | 0 | 4 | 1 | 2 | 4 | 30 |
|  | positive | 2 | 6 | 4 | 5 | 8 | 5 | 8 | 7 | 4 | 49 |
|  | zero | 0 | 0 | 0 | 0 | 1 | 0 | 0 | 0 | 1 | 2 |
| Total |  | 9 | 9 | 9 | 9 | 9 | 9 | 9 | 9 | 9 | 81 |
| **b** |  |  |  |  |  |  |  |  |  |  |  |
| Count |  |  |  |  |  |  |  |  |  |  |  |
|  |  | Taxon |  |  |  |  |  |  |  |  |  |
|  |  | Betula | Bistorta | Campanula | Cerastium | Draba | Equisetum | Graminoids | Salix | Stellaria | Total |
| Abundance response ratio category | missing | 0 | 3 | 4 | 3 | 8 | 0 | 0 | 0 | 1 | 19 |
|  | negative | 0 | 5 | 2 | 5 | 1 | 8 | 8 | 9 | 7 | 45 |
|  | positive | 9 | 0 | 2 | 1 | 0 | 1 | 1 | 0 | 1 | 15 |
|  | zero | 0 | 1 | 1 | 0 | 0 | 0 | 0 | 0 | 0 | 2 |
| Total |  | 9 | 9 | 9 | 9 | 9 | 9 | 9 | 9 | 9 | 81 |

